# Mutual predators: A descriptive cross-sectional study to identify prevalence and co-relation of Hepatitis C Virus and Human Immunodeficiency Virus type-1 coinfection

**DOI:** 10.1101/017574

**Authors:** Fouzia Ashraf, Dalaq Aiysha, Muhammad Tajamal, Shahzeb Javed, Saamia Tahir, Omar Ali, Mahmood Shaukat

## Abstract

**Background:** Coinfection, bacterial or viral origin, in HIV infected individuals’ remains to be the only leading cause of deaths. This study was designed to analyze received plasma samples and plasma samples of referred patients for HIV testing to detect HIV and HCV mono and co-infection by real time PCR and finding co-relation of viral load of both viruses. Highlight and magnify the hidden coinfection, prior to seroconversion, of HIV type-1 and Hepatitis C Virus in received samples.

**Methods:** Analyses were based on randomly selected 78 patients’ stored plasmas. Plasma samples were tested for both, HIV-type 1 and HCV viral RNA by real time PCR. Statistical formulas were used to identify men and the inter quartile range of patients age. The data were analyzed by IBM SPSS Statistics 21 (SPSS Inc., Chicago, IL). Study variables include gender, age and viral loads of HIV type-1 and HCV. Pearson correlation was used to evaluate any correlation in study variables.

**Result:** Prevalence of HCV was 10.3%, HIV-type 1 was 19.2% and their co-infection was 37.2 percent. Thirty three percent individuals had no infection of both viruses. Gender based distribution showed that 74.4% (58/78) sample population was male. The mono-infection and co-infection was higher in males (39.7%) and highest viral load too. There was a positive correlation (CI= 95%) between the two variables; HIV and HCV viral loads, as r = 0.736, n=29, p= 0.001.

**Conclusion:** Prevalence of HIV type-1 and HCV mono-infection and co-infection was higher among males as compared to females. Increased viral load was also evident among male co-infected individuals. This study proved the emergence of HCV coinfection in HIV infected individuals, and a need for on time diagnosis and treatment.

## Introduction

Target cells of both viruses-Human Immunodeficiency Virus and Hepatitis C Virus-are different, but in co-infection their outcomes are deleterious due to increased disease progression in combined proliferation (Kim & Chung, 2009). Common mode of HIV and HCV transmission is by direct contact with body fluids of an infected person (Polsky, Kim & Chung, 2000–2001). Globally, prevalence of viral hepatitis among HIV infected people is seven million worldwide (Soriano et al., 2010). The prevalence rate of co-infection varies considerably between and within countries, depending on rate of risk factors in the population. Viral hepatitis is an emerging non-AIDS related, cause of morbidity and mortality among HIV patients (Bica et al., 2001).

In HCV mono-infection about fifteen to forty five percent patients with acute viral infection clear the virus, while twenty to fifty percent develop persistent viremia (Thomas & Seeff, 2005). The persistence of HIV and HCV is attributed to high rates of replication, its ∼10^12^ and ∼10^9^ virion/day for HCV and HIV, respectively (Kim & Chung, 2009). Transmission of communicable disease has been an influential element in acquiring infection and disease progression. The highest rate of HIV type-1 and HCV co-infection has been reported to be 90 percent among in-vitro drug users (Sherman et al., 2002). Hemophiliacs are the second dominant group of co-infection, who received contaminated blood or its products (Yee et al., 2000). The reported prevalence of HIV and HCV among blood donors of Pakistan is was, 3.78 and 0.06 percent respectively (Sultan, Mehmood &Mahmood, 2007; Waheed et al., 2009; Waheed et al., 2010) Co-transmission of HIV and HCV can occur through percutaneous route among injection drug users, but they were usually infected first with HCV (Di Martino et al., 2001).

Hepatitis C Virus and HIV type-1 both belongs to two different families of viruses. This particular association has a major effect on their life cycle. HCV replicates through RNA-dependent RNA polymerase, (Bartenschlager, Lohmann & Lohmann, 2000) while HIV type-1 first integrates its genetic material in host DNA through the viral reverse transcriptase enzyme and then replicates (Zheng, Lovsin &Peterlin, 2005). There is a direct relation to inoculums size and acquiring HCV infection, after transmission through blood transfusion or by contaminated needles, viremia occurs within days or it takes six to eight weeks, respectively (Maheshwari, Ray & Thuluvath, 2008). In America, a large cross-sectional analysis (n=1687) of two HIV trials, reported that 75% subjects had HCV RNA level above 800,000 IU/ml (Thomas & Seeff, 2005).

## Materials and Methods

### Sample sources and preparation

A cross-sectional study was conducted during the months of May, June and July 2014, among referred patients to Jinnah Hospital Lahore (JHL). JHL is the second largest teaching hospital, which provides health care facilities to the inhabitants of Lahore and adjacent areas. A random blood samples were collected from patients along with their age and gender. Laboratory testing was performed in molecular diagnostic lab, specialized for PCR testing of infectious diseases. Plasma from each sample was separated by centrifugation and stored at -20^°^C until analyzed.

### Nucleic acid extraction and amplification

Viral RNA was extracted from 200μl of plasma using QIAamp VIRAL RNA mini-kit from Qiagen (CAT. No. 52904) from Germany. The PCR quantification of samples was carried out by Artus RT-PCR kit for HIV and HCV by Qiagen, for all randomly selected samples, regardless of their serological result. Standard procedures, as proposed by Kwok and Higushi [1989], were followed to avoid contamination.

### Study design and statistical analysis

All study subjects were distributed in three groups, according to their age as, minors (< 18 years), adult (>18 and < 50 years) and old (>50 years). They were also distributed according to gender; in three groups as follows: male, female and transgender. The mean age and inter quartile range were also calculated by statistical formulas. All data were analyzed by IBM SPSS Statistics 21 (SPSS Inc., Chicago, IL). Study variables include gender, age and viral loads of HIV type-1 and HCV.

## Results and Discussion

The overall prevalence of co-infection, HIV and HCV mono-infection among walk-in patients of Jinnah Hospital, was 29 (37.2%), 15 (19.2%) and 8 (10.3%) respectively. Among 78 total participants, 26 (33.3 %) had no infection of HIV and HCV. Of the study population (n=78), 25.6% (20/78); 74.4% (58/78) were female and males respectively. The mean age of the sample population, irrespective of gender, was 30 years with an inter-quartile range of 24 to 40. The youngest positive co-infected person was male of seven years and the oldest one male of 59 years.

As shown in table 1 and figure 2; among 57 male participants, 23 (39.7%) had HIV and HCV co-infection, 13 (22.4%) carried the HIV mono - infection, 5 (8.6%) carried the HCV mono - infection and rest 17 (29.3%) had no infection of both viruses. Among twenty female, the distribution of HIV or HCV mono-infection and co-infection of both (HIV and HCV) was 2 (10%), 3 (15%) and 6 (30%), respectively. Nine females, representing 45% of the female population (n=20), had no infection of both HIV and HCV. There was a positive correlation between the two variables; HIV and HCV viral loads, as r = 0.736, n=29, p= 0.001. Figure 1 shows the prevalence of HIV and HCV co-infection, HIV type 1 and HCV mono-infection in the study population, while table 2 shows the number of cases of HIV and HCV co-infection, mono-infection and no infection of both viruses in different age groups.

**Figure 1:**
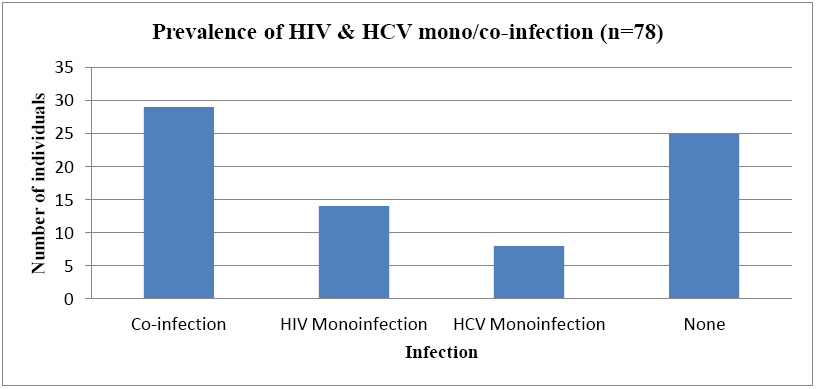
Prevalence of HIV and HCV mono or co-infection in sample population (n=78)

**Table 1.**
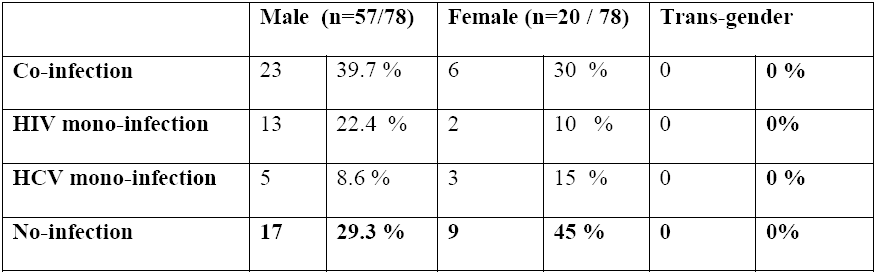
Prevalence of HIV and HCV in different gender groups.

**Figure 2:**
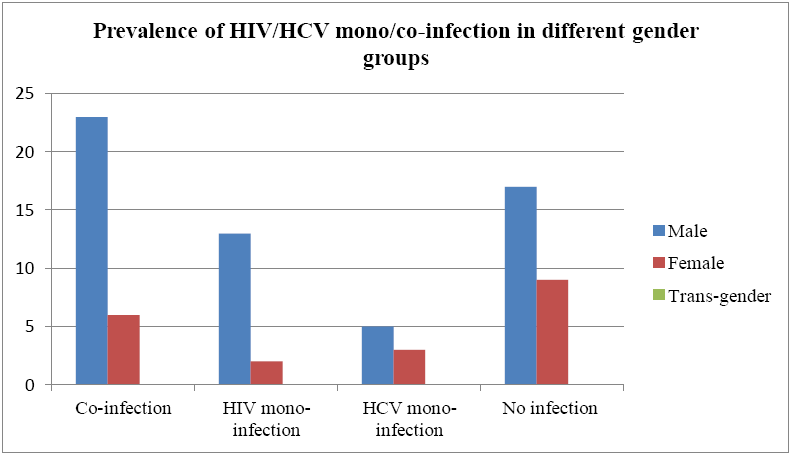
Prevalence of HIV and HCV mono or co-infection in different gender groups

**Table 2.** Prevalence of HIV and HCV in different age groups.

|  | **Minor $\leq 18$ years** |  | **Adult $>18$ & $< 50$ years** |  | **Old $\geq 50$ years** |  |
| --- | --- | --- | --- | --- | --- | --- |
| **Co-infection** | 3 | 3.8% | 25 | 32.1 % | 1 | 1.3 % |
| **HIV mono-infection** | 0 | 0 % | 11 | 14.1 % | 3 | 3.8% |
| **HCV mono-infection** | 0 | 0% | 6 | 7.7 % | 2 | 2.6 % |
| **No-infection** | 1 | 1.3 % | 23 | 29.5 % | 3 | 3.8% |

The results we present here show the prevalence of HIV type-1 and HCV co-infection among walk-in patients of Lahore region. The analysis revealed increased prevalence of co-infection of both persistent viruses. An estimated five to ten million people in the world are living with HIV and HCV co-infection (Operskalski & Kovacs, 2011). The detection of HCV RNA, during initial HCV infection, is possible after two to fourteen days of exposure (Hajarizadeh, Grebely J & Dore, 2013). The seroconversion of Hepatitis C Virus is delayed in HIV positive co-infected individuals, possibly due to the immunosupression trade mark of HIV infection. Only three studies revealed zero to thirteen percent incidence of seronegative chronic Hepatitis C virus infection (Thio et al., 2000; Chamie et al., 2007; Thomson et al., 2009). Such silent and persistent infection not only alters the course of HIV disease progression, but also results an ineffective treatment and its outcomes (Antonucci et al., 2005; Hua et al., 2013). Therefore, on time and accurate diagnosis is vital for survival.

This study established the presence of HIV and HCV co-infection among general, random infected population of Punjab. Among the HIV positive individuals (n=43) of our study, 67.5 % were co-infected with Hepatitis C Virus. The highest (90%) reported HIV and HCV co-infection was among jail inmates of Sindh, Pakistan (Safdar, Mehmood & Abbas, 2009); followed by 73.74% in Lahore inmates (Nafees et al,, 2011). The highest prevalence of co-infection; 32.1 % has been seen among the adult population of our study. The prevalence of HCV co-infection was less in our study as compared to the reported prevalence; 53.54% of Nafees et al., (2011).

In the sample population, no significant statistical relation was found between age and gender groups with an incidence of mono or co-infections. However, in our study, the male population (74%) as compared to the female population (26%) had a higher prevalence of HIV and HCV mono/co-infections. Our study has a higher percentage of male population as compared to other recently reported study (Khan, Ali & Awan, 2013), but less as compared to the study of jail inmates in Lahore (Nafees et al., 2011). Three minor individuals in our study; aged 7, 15 and 17 years, had co-infection of HIV and HCV; among them one was female of 15 years. A prevalence of coinfection in minor population (3.8 %) was observed in our data which was not reported before in any other studies (Nafees et al., 2011; Khan, Ali & Awan, 2013). The mean age of our study group was 30 years old. This mean age was equal to the sample population of previous reported study (Samo et al., 2013) but, more as compared to other reported studies from this region irrespective of their risk population and gender (Nafees et al., 2011; Khan, Ali & Awan, 2013).

This study was aimed to see any kind of correlation between viral load of HIV type-1 and HCV co-infected individuals. Pearson’s correlation model showed evidence of significant (P<0.001) supporting results of our hypothesis (CI=95%). Positive and less than one value of r (r = 0.736), indicated strong relationship between viral load and correlation in pathogenesis of HIV type-1 and HCV. This correlation indicated that increase in HIV type-1 viral load associated with an increase in HCV viral load. Many other studies have shown similar results of correlation between HIV type-1 viral load an HCV viral load (Sherman et al., 1993; Mazza et al., 1994; Eyster et al., 1994; Cribier et al., 1995; Thomas et al., 1996; Cribier et al., 1997; Beld et al., 1998). Immediate prevention and surveillance measures are needed to control the spread of mono-transmission as well as co-transmission of HIV type-1 and HCV infections.

## Conclusions

The prevalence rate of coinfection of HIV type-1 and Hepatitis C virus is much higher in our study conducted in a metropolitan city, Lahore, Pakistan. Co-infection of Hepatitis C Virus is associated with increase viral replication helping in disease progression of both viruses, HIV type-1 and HCV, irrespective of gender and age. Health care professionals should include molecular diagnosis of HCV during a routine evaluation of HIV suspected and infected individuals.

**Figure 3:**
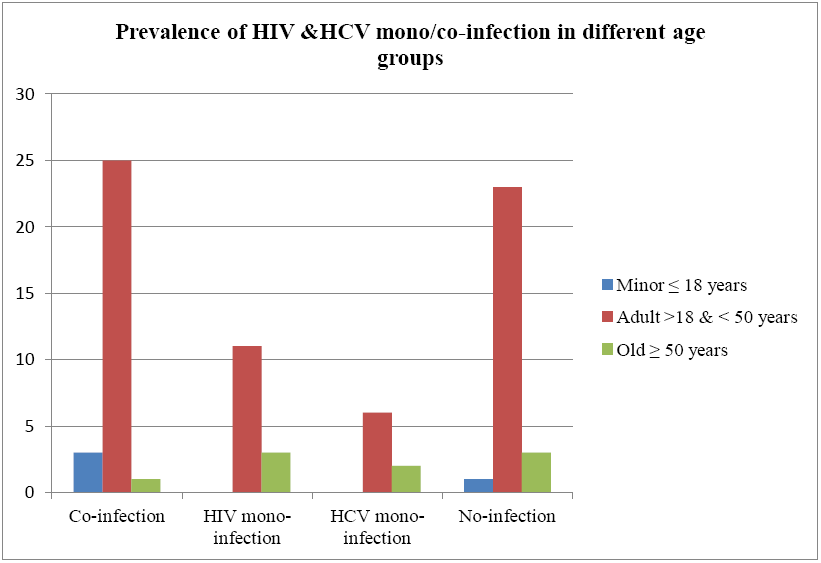
Prevalence of HIV and HCV in different age groups (n=78)

## Acknowledgement

This study was not possible without our sample sources and laboratory staff.

